# Evaluation of cerebellar function scores in relation to cerebellar axonal loss in multiple sclerosis

**DOI:** 10.1101/2020.05.15.094938

**Authors:** F. Boonstra, S. Gajamange, G. Noffs, T. Perera, M. Strik, A. Vogel, H. Butzkueven, A. Evans, A. van der Walt, S. Kolbe

## Abstract

**Background:** Cerebellar damage is common in people with multiple sclerosis (pwMS) and is associated with worse progression and relapse recovery. Studies into the importance of the cerebellum in pwMS are hampered by limited understanding of cerebellar damage and its relation to cerebellar function in pwMS.

**Objective:** Examine axonal loss, as a primary driver of progressive neurological decline, in the cerebellum using advanced diffusion MRI and compare axonal loss with cerebellar dysfunction in pwMS

**Methods:** We recruited 55 pwMS and 14 healthy controls. Clinical assessments included scale for the assessment and rating of ataxia (SARA), and Bain tremor ratings. Subjects underwent FLAIR, T1-weighted and diffusion MRI. Cerebellar grey and white matter and lesion volume were calculated. Cerebellar axonal loss was examined with fibre-specific markers. Fibre density and cross-section (FDC) accounts for microscopic and macroscopic changes in a fibre bundle.

**Results:** Loss of cerebellar FDC was associated with increased SARA (r=-0.42, p<0.01) and tremor severity (rho=-0.35, p=0.01). Cerebellar lesion volume correlated with SARA (r=0.49, p<0.01) and tremor severity (rho=0.41, p=0.01).

**Conclusion:** Fibre-specific measures of cerebellar pathology could provide a functionally relevant marker of cerebellar damage in MS. Future trials using fibre-specific markers are needed to further characterize cerebellar pathology in pwMS and understand its significance in disease progression.

## Introduction

Multiple sclerosis (MS) is a chronic inflammatory and neurodegenerative disease of the central nervous system. The clinical course of MS is highly heterogenous and outcomes are difficult to predict. The cerebellum is an important structure deeply integrated into major networks involved in motor control, cognition and executive control (1). In MS, focal damage to cerebellum and its efferent and afferent pathways lead to a range of highly disabling symptoms such as loss of dexterity, ataxia, dysarthria and tremor (2) that have a significant impact on quality of life (3, 4). Importantly, involvement of the cerebellum at disease onset is associated with worse prognosis (5) and, cerebellar relapses are associated with poorer recovery and earlier onset of progressive disease (6). To improve our knowledge about the role the cerebellum plays in MS, it is important to understand cerebellar damage and its relation to cerebellar function.

Cerebellar inflammation and neurodegeneration in people with MS are commonly identified using magnetic resonance imaging (MRI). Lesions in the white matter, including the cerebellar peduncles (7), and cerebellar atrophy (8) are common and more severe in patients with progressive disease phenotypes. Despite the importance of cerebellar involvement to general disease progression, MRI indicated cerebellar pathology is often not reliably detectable in patients in the early stages of MS (9). Furthermore, it correlation with clinical disease severity and cerebellar function, measured using EDSS and cerebellar functional system score (cFSS) respectively, is only modest at best (10, 11). This likely reflects a combination of pathologically non-specific imaging markers and complex interactions between focal pathology, and individual disability trajectories. Thus, the drive towards personalised treatment requires greater specificity and sensitivity in terms of both pathological and disability markers.

Despite the constellation of pathologies present in the MS brain, irreversible axonal loss is considered to the primary contributor to progressive neurological decline (12-14). Axons are tightly bundled and due to the strong anisotropy of water diffusion within the axolemma, it can be readily interrogated using diffusion-weighted MRI (dwMRI). The cerebellum receives nearly 200 million input fibres (15) and is known to consist of complex networks with large number of crossing fibres (16, 17). Advanced dwMRI methods can assess fibre-specific changes in the presence of crossing-fibres and cellular pathologies such as inflammation, demyelination and oedema (18). We have shown such methods are highly sensitive to axonal loss in people with MS presenting with optic neuritis, revealing with a high specificity, damage to the visual pathways of the brain relevant to the specific neurological presentation of patients (19).

In this study we aimed to assess the relationship between the clinical presentation of cerebellar dysfunction and cerebellar pathology using advanced dwMRI and conventional MRI techniques (brain and lesion volumetrics) in MS. Cerebellar function was assessed using a range of clinical assessment tools, including cerebellar functional system score (cFSS), scale for the assessment and rating of ataxia (SARA), and Bain. Determining which measure of cerebellar function shows the strongest relation to cerebellar pathology, could inform the most adequate measure to monitor cerebellar function in the clinic.

## Methods

### Study design

Patients were recruited from a large ongoing randomized, controlled trial of the efficacy and safety of BoNT-A in MS-related arm tremor (ACTRN12617000379314). The study was approved by the Melbourne Health Human Research Ethics Committee. All participants gave informed voluntary written consent.

### Participants

We recruited 55 people with MS (pwMS) and 14 healthy controls. Detailed medical and cerebellar dysfunction history were obtained for all participants.

### Clinical assessment

A Neurostatus certified neurologist determined the Kurtzke Expanded Disability Status Scale (EDSS) for all participants as a measure of overall disease disability (20). Three different scores were measured for cerebellar dysfunction: cFSS, the scale for the assessment and rating of ataxia (SARA) (21) and the Bain score (22). The SARA is a validated MS-specific ataxia scale and thought to provide a comprehensive assessment of cerebellar function. The SARA is a (semi-) quantitative 40-point scale rating gait, stance, sitting, speech disturbance, finger chase, nose-finger test, fast alternating hand movements, and heel-shin slide. The Bain score was used to rate tremor severity observed while writing a standardised sentence (‘This is a sample of my best handwriting’) and drawing an Archimedes spiral on a pre-drawn pattern with the dominant hand (22, 23). The three Bain scores are rated from 0-10 with 0 meaning no tremor and 10 extremely severe tremor. Bain total was calculated as the summation of the three different scores.

### Imaging protocols

Participants were imaged using a 3T MRI system (Trio, Siemens, Erlangen). Structural imaging included 1) high-resolution three-dimensional (3D) T1-weighted magnetization-prepared rapid acquisition with gradient echo (MPRAGE) scan with prospective motion correction (recovery time = 2530 ms; echo time = 2.5 ms; inversion time = 1260 ms; field of view = 176 × 256 mm; voxel size = 1.0 × 1.0 × 1.0 mm), 2) high-resolution 3D T2-weighted double inversion recovery (DIR) sequence (repetition time = 7400 ms; echo train = 324 ms; inversion time = 3000 ms; flip angle = 120°; echo train length = 625; field of view = 144 × 220 mm; voxel size = 1.0 × 1.0 × 1.0 mm) and (3) multi-shell dwMRI (voxel size = 2.4 x 2.4 x 2.4 mm, 16 non-diffusion images, 31 b = 1000 s/mm^2^, 51 b = 2000 s/mm^2^, 64 b = 2800 s/mm^2^).

### Volumetric analyses

T1-weighted MRI was analysed using FreeSurfer version 6.0 (http://surfer.nmr.mgh.harvard.edu/) and Osirix Lite v9.0.2. Freesurfer was used to calculate whole brain volume, cerebellar volume using the standard recon-all pipeline (24). Lesions were identified on DIR images using the lesion prediction algorithm within Lesion Segmentation Toolbox (LST) version 2.0.15 for SPM version 12 (25). To optimize the automatic lesion segmentation, we manually determined the optimal threshold for each participant and manually corrected the final lesion maps. Lesion volume within the cerebellum was calculated by the intersection between FreeSurfer region of interest mask and lesion masks.

### Diffusion analyses

Diffusion imaging data were pre-processed and analysed with MRtrix3 (http://www.mrtrix.org) and the multi-shell multi-tissue fixel based analysis pipeline (described in detail at https://mrtrix.readthedocs.io/en/latest/fixel_based_analysis/mt_fibre_density_cross-section.html). Briefly, dwMRI data were denoised, corrected for eddy current and susceptibility induced distortions, up-sampled by a factor of two, and multi-shell multi-tissue constrained spherical deconvolution (MSMT-CSD) was used to calculate the continuous fibre orientation distribution (FOD) for each voxel (26). A population-average FOD template was created from all subject specific FOD images using an iterative co-registration approach (18). Each FOD within the FOD template was segmented to identify individual fibre populations (termed “fixels”), resulting in a fixel template mask. Lastly, each subject’s FOD map was nonlinearly aligned to the FOD template and segmented to identify subject-wise fixels. Fixels were reorientated and correspondence was calculated between the subject-wise fixels and the template fixels, allowing for fixel-wise univariate statistical analyses. Registrations were visually checked for errors with no gross errors observed.

For each fixel, the modulated fibre density was calculated as the multiplication of the volume integral of the FOD lobe [termed fibre density or FD, previously shown to be proportional to the intra-axonal volume of the fibre population (18, 27)] and the deformation warp field perpendicular to the fixel generated from the registration to the template (termed fibre cross-section of FC, accounting for differences in cross-sectional area of white matter fibre bundle). Thus the modulated fibre density (FDC) can be considered as a marker of the total number of axons in the tract after accounting for variation in both fibre density (FD) and fibre cross-section (FC) (18). Previous studies have reported results from FD, FC and FDC [for example (19, 28)]. Here we only report FDC, as, (1) in-house investigations have shown that FD is often reduced in the edges of atrophic tracts and therefore the correction for FC is important for accounting for subtle inconsistencies in spatial alignment, and (2) volumetric changes were measured using high resolution T1-weighted scans.

A pos-hoc tract-level analysis was performed to examine FDC measures within the cerebellum. A mask of the cerebellum was applied. To examine group differences in cerebellar FDC between pwMS and controls, and correlations between FDC and functional variables in patients, we used fixel-wise univariate tests corrected for multiple comparisons using connectivity-based fixel enhancement (29). All tests were corrected for age and sex and used 5000 permutations for calculating null test distributions. Finally, for each participant we extracted the average FDC of all cerebellar fixels to be used in subsequent statistical analyses.

### Statistical analyses

All analyses were performed in SPSS version 25. Tests for normality showed that EDSS, cFSS, Bain scores and lesion load variables were non-normally distributed, while the SARA, FDC and cerebellar volume were normally distributed. Log 10 transformation improved normality for lesion load measures. As log transformation did not improve normality for cFSS and Bain scores, nonparametric test were used.

Demographics, volumetric measures and cerebellar FDC were compared between pwMS and controls using independent t-tests. Cerebellar volume and lesion volume were corrected for differences in intracranial volume.

For parametric analyses, a partial Pearson correlation analyses was performed between volumetric measures (FDC, cerebellar volume and lesion volume) and the clinical measures (SARA, cFSS and Bain scores) corrected for age and gender. For non-parametric analyses, we used linear regression to correct for age and gender followed by Spearman correlation tests.

## Results

Summary of all demographic, clinical and imaging variables for both healthy and pwMS is depicted in Table 1.

**Table 1.** Demographics, disease characteristics and imaging measures.

|  |  | Healthy controls<br>(n=14) | MS participants<br>(n=55) | Test (p) |
| --- | --- | --- | --- | --- |
| Disease and Demographics | Age, mean (sd) | 45.29<br>(15.7) | 46.80 (12.1) | t = -0.39,<br>p = 0.695 |
|  | Gender, female % | 57.1 | 78.2 | t = 1.42,<br>p = 0.173 |
|  | Disease duration, mean (sd) | - | 12.8 (7.9) | - |
|  | Disease course, progressive % | - | 54.5 | - |
| Clinical and Cerebellar Function | EDSS, median (IQR) | - | 3.5 (2,5.5) | - |
|  | cFSS, median (IQR) | - | 2 (1,2) | - |
|  | SARA, mean (sd) | - | 8.27 (6.0) | - |
|  | Bain severity, median (IQR) | - | 2 (0,3) | - |
|  | Bain handwriting, median (IQR) | - | 1 (0,3) | - |
|  | Bain Archimedes, median (IQR) | - | 1 (0,3) | - |
| Lesion Load | Cerebellar lesions, mean (sd) | - | 39.5 (75.0) | - |
| Cerebellar Volumetrics | Cerebellar white matter x10<sup>4</sup>, mean (sd) | 2.79 (0.40) | 2.62 (0.36) | t = 1.52,<br>p = 0.133 |
|  | Cerebellar grey matter x10<sup>4</sup>, mean (sd) | 11.0 (1.67) | 10.3 (1.07) | t = 1.37 p = 0.191 |
| Cerebellar diffusion | FDC cerebellum, mean (sd) | 0.26 (0.03) | 0.25 (0.03) | **t = 2.16,**<br>**p = 0.035** |

**Table 2.**
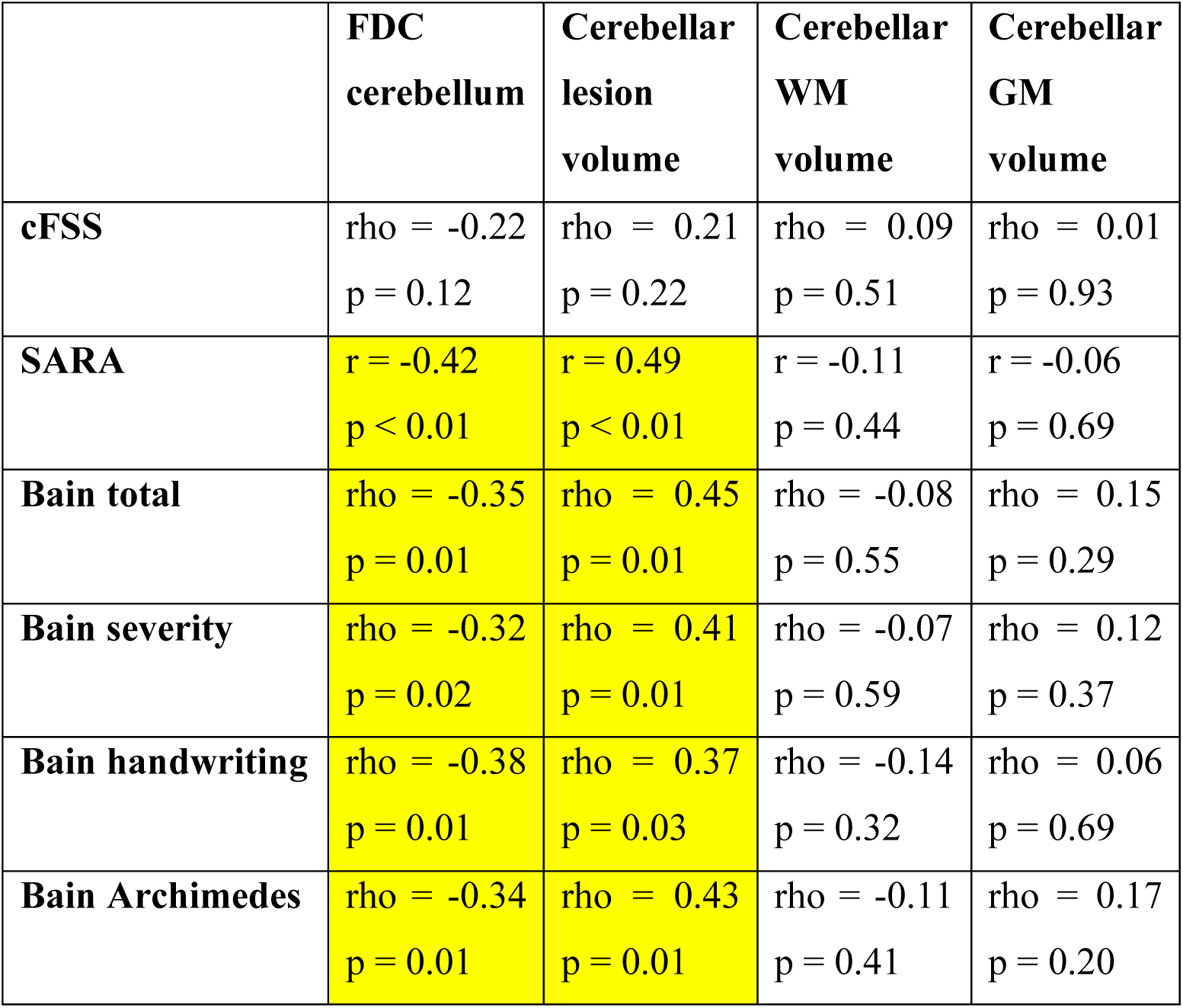
Correlation between disease characteristics and imaging measures of the cerebellum.

Cerebellar FDC was compared between subject groups. We found that pwMS [0.25 (0.03)] had significantly lower cerebellar FDC than healthy controls [0.26 (0.03)] (t = 2.16, p = 0.035). No significant differences were found in cerebellar FDC between patients with relapsing-remitting and secondary progressive MS.

### Correlation analyses

We observed significant negative correlation between the SARA score and cerebellar FDC (r = −0.42, p < 0.01). Furthermore, all upper limb tremor severity scores showed a significant moderate negative correlation with cerebellar FDC (Bain total rho = −0.35, p = 0.01, Bain tremor severity rho = −0.32, p = 0.02; Bain handwriting rho = −0.38, p < 0.01 and Bain Archimedes spiral rho = −0.34, p = 0.01). See Figure 1 for a visual representation of the correlation between FDC and clinical cerebellar function.

**Figure 1.**
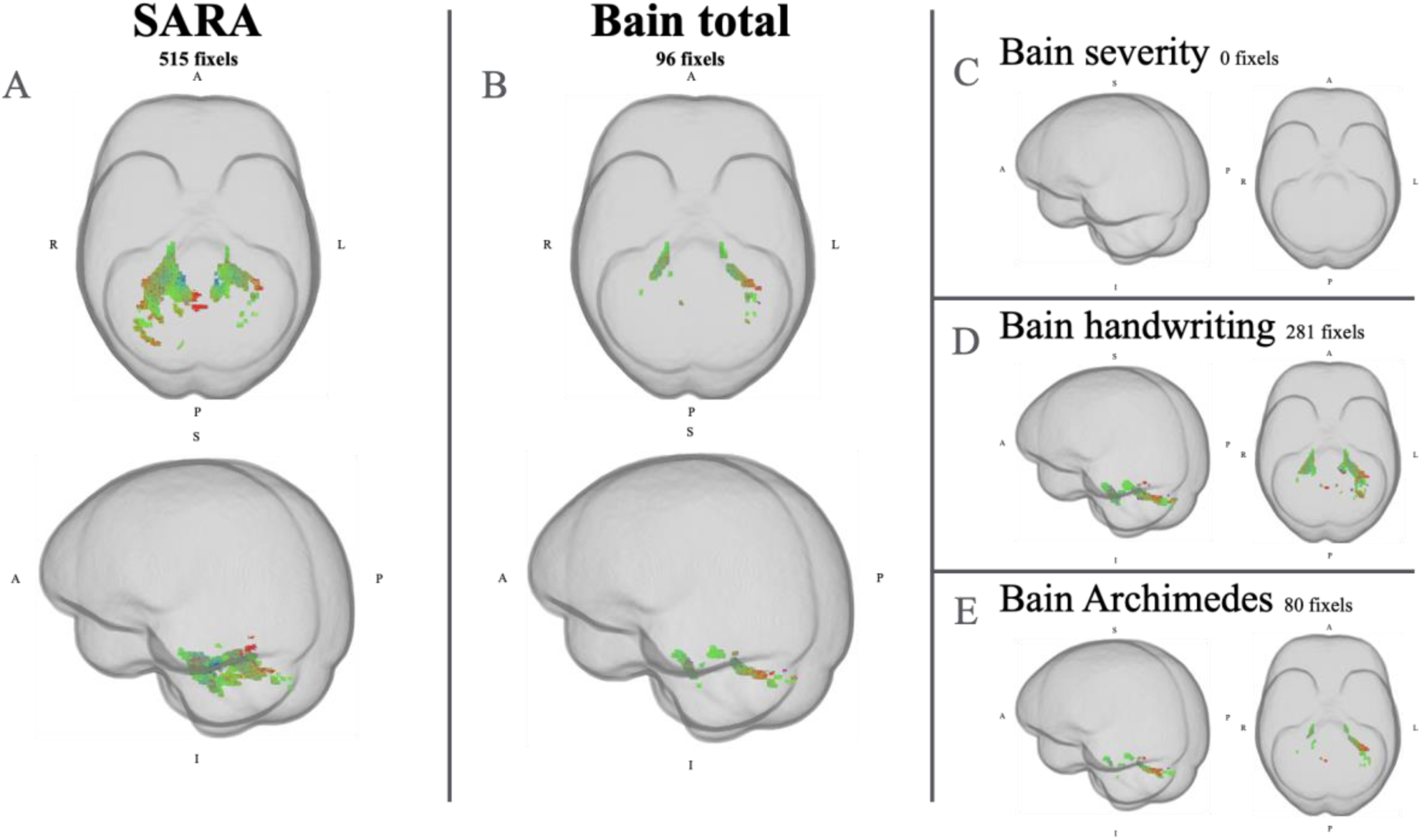
Fixels showing significant correlations between cerebellar FDC and cerebellar dysfunction: A) SARA, B) Bain total, C) Bain severity, D) handwriting and E) Archimedes (right). Total number of fixels significant (threshold at *p*_*fwecorr*_<0.05) depicted. Fibre colours are directional: blue = superior-inferior direction; red = left-right direction; green = anterior-posterior direction.

Cerebellar white and grey matter volume did not correlate with any of the clinical measures. Cerebellar lesion load significantly correlated to SARA (rho = 0.49, p < 0.01), and all upper limb tremor severity scores (Bain total rho = −0.45, p = 0.01, Bain tremor severity rho = −0.41, p = 0.01; Bain handwriting rho = −0.37, p = 0.03 and Bain Archimedes rho = −0.43, p = 0.01).

The post-hoc correlations between the cerebellar FDC and clinical measures of cerebellar function are displayed in figure 1. The SARA and Bain scores all showed areas of significant correlation with the FDC. Specifically, the SARA correlated with the largest region within the cerebellum (515 fixels at *p*_*fwecorr*_<0.05), followed by Bain handwriting (281 fixels), Bain total (96 fixels), Bain Archimedes (80 fixels) and Bain severity (0 fixels).

### Cerebellar axonal damage and conventional MRI measures

Cerebellar FDC significantly correlated with cerebellar white matter volume (r = 0.25, p = 0.04) and cerebellar lesion load (r = −0.33, p = 0.01). No correlation was found between the cerebellar FDC and cerebellar grey matter. These results are reported in supplementary table 1

## Discussion

This study aimed to examine the relationship between clinical measures of cerebellar function and cerebellar axonal loss in pwMS. We found significant reduction in cerebellar FDC in pwMS compared to healthy controls. FDC is the apparent fibre density modulated by the jacobian of the nonlinear transformation to the study template perpendicular to the tract. The modulation accounts for differences in cross-sectional area of the tract that could influence axonal density. Such differences could relate to individual differences in healthy anatomy or pathological atrophy. Thus FDC is marker for the total axonal number in a tract (27). Previous neuroimaging studies have also reported fibre density (FD) and fibre cross-section (FC) (19, 28), however, here we chose to focus on FDC exclusively, firstly to ensure FD was not confounded by variability in FC (nor FC confounded by FD), and secondly, to reduce the number of statistical tests performed. In this study we focused exclusively on the cerebellum for all fixel-wise statistical analyses. While it is well known that white matter pathways to and from the cerebellum include tracts outside the anatomical boundary of the cerebellum “proper”, we chose to focus our analysis in this way for simplicity and because pathology to connected tracts outside the cerebellum ROI would still affect cerebellar tracts within the cerebellum due to Wallerian or retrograde axonal degeneration.

This is the first study to apply fixel-wise analysis of advanced diffusion imaging to the cerebellum in pwMS, however, previous studies have examined cerebellar diffusion tensor imaging (DTI) in the context of disease stage and specific disabilities such as mobility loss and cognition. While care needs to be taken when interpreting cerebellar DTI results, as they do not adequately model multiple fibre population (i.e. crossing fibres), CSF and other non-axonal cell types (30-32), the results are nonetheless important for contextualising the current study. Anderson *et al.* (33) studied RRMS, PPMS and control groups reported no difference in MCP or SCP DTI in RRMS compared to controls, and significant DTI changes in PPMS compared to both RRMS and controls that correlated with time to complete a 25-foot walk. It is conceivable that the lack of significant difference in the RRMS group could be due to low sensitivity due to low number of subjects studied (RRMS, n=12; PPMS, n=11). In contrast, Deppe and colleagues (9) noted in a larger group of patients (n=68), significant loss of DTI fractional anisotropy (FA, a marker of reduced axonal density and demyelination) in the absence of significant atrophy in patients with early MS (disease duration < 50 months), while patients with later stage MS (mean disease duration 113 months) exhibited both significant loss of both FA and atrophy. Interestingly, there was no difference in FA between the early and late MS groups suggesting that cerebellar axonal loss manifests early in the disease as loss of axonal density (loss of FA) with preserved volume, while in later stages, atrophy emerges with preserved loss of axonal density. These findings support the need for measures of axonal density and tract atrophy to be considered in parallel in order to fully elucidate axonal loss in a structure, as is the case for the fixel-based analysis of FDC performed here. Moroso *et al.* (34) explored relationships between cognitive functions and cerebellar DTI parameters measured from specific cerebellar regions, reporting several cerebellar regions where loss of FA correlated significantly with working memory and information processing speed. Together these studies highlight the importance of cerebellar damage to the progression of a range of disabilities common in pwMS. Kolbe *et al.* (35) also noted significant correlations between cognitive function and DTI changes in cerebellar tracts and atrophy of specific cerebellar lobes in patients with relapsing-remitting MS.

We found that clinical measures of cerebellar dysfunction are closely related to cerebellar damage indicated by fixel based analysis. This suggest that pwMS with more extensive axonal loss within specific cerebellar regions have worse cerebellar function, measured using the SARA and the Bain scores. The former showing the strongest and most extensive association with cerebellar axonal damage. To our knowledge, to date no other studies have looked at the relationship between axonal damage and measures of clinical cerebellar function in MS. In patients with autosomal dominant spinocerebellar ataxia, Adanyeguh et al. (36) found a negative correlation between the SARA score and FDC along the corticospinal tract. Two studies using diffusion tensor imaging of the cerebellum in ataxia patients (spinocerebellar ataxia type 10 and Friedreich ataxia), found a very weak to no correlation with SARA (37, 38).

This could reflect the inability of diffusion tensor imaging to accurately capture cerebellar diffusion. Together, our findings suggest that axonal loss is measured using fibre-based analyses capable of understanding the relationship between cerebellar damage and cerebellar dysfunction in MS.

No gold-standard validated tool for measuring MS-cerebellar function exists. Moves to improve the characterisation of the clinical presentation of cerebellar damage in MS are underway (39). The main clinical measure of MS disability (EDSS) (20) offers only rough estimates of cerebellar function and it is not a very sensitive measure of change (40). This study showed that both the SARA and the Bain scores are moderately correlated with cerebellar damage in MS, with the SARA showing the strongest correlation with both axonal loss and cerebellar lesion load. These results suggest that the SARA and Bain might be a better clinical measure of cerebellar function in MS compared to EDSS. Further research into the sensitivity of the SARA and Bain is needed, to determine if it can be used as an adequate tool for monitoring cerebellar function in MS.

This is only the second study to have applied fixel-based analyses in MS. The first study used a sample of early pwMS with minimal pathology to the brain (19). Pathologically severe lesions with complete destruction of the fibre bundle can potentially affect image registration locally. The present study used a sample of pwMS with a more heterogeneous disease, including higher disease severity and more extensive brain pathology. During our analyses we visually checked image registration and did not identify any errors. Even if small differences in registration had occurred, these would have mostly been accounted for by using FDC rather than fibre density and cross-section separately. In addition, the use of the combined measure also allows for a less biased and more sensitive measure compared to fibre density and cross-section separately as it does not suffer from inherent inter-dependency (18). More studies are needed to study the effect lesions have on the fixel-based analyses. Furthermore, future longitudinal studies using larger samples could help to understand how axonal damage evolves over time.

To conclude, we showed significant cerebellar axonal damage in pwMS. Furthermore, cerebellar pathology is associated with cerebellar function in MS as measured using the SARA and to a lesser extent Bain. These findings both help our understanding of cerebellar pathology in MS and suggest that SARA could be used as a clinical tool to monitor cerebellar damage.

---

**Supplementary table 1.** Correlation between fixel-based analyses and conventional MRI measures of the cerebellum.

|  | **FDC cerebellum** |
| --- | --- |
| **Cerebellar white matter** | $r = 0.25, p = 0.04$ |
| **Cerebellar grey matter volume** | $r = 0.14, p = 0.25$ |
| **Cerebellar lesions** | $r = -0.33, p = 0.01$ |

